# Interpreting mosquito feeding patterns in Australia through an ecological lens; an analysis of blood meal studies

**DOI:** 10.1101/492934

**Authors:** E.B. Stephenson, A. Murphy, C.C. Jansen, A.J. Peel, H. McCallum

**Affiliations:** Environmental Futures Research Institute, Griffith University, Queensland; QIMR, Herston, Queensland, 4006, Australia; Communicable Diseases Branch, Department of Health, Queensland Government, Herston, Queensland, 4006

**Keywords:** Blood meal, associations, vector, vertebrate, disease risk, blood feeding

## Abstract

Mosquito-borne pathogens contribute significantly to the global burden of disease, infecting millions of people each year. Mosquito feeding is critical to the transmission dynamics of pathogens, and thus it is important to understanding and interpreting mosquito feeding patterns. In this paper we explore mosquito feeding patterns and their implications for disease ecology through a meta-analysis of published blood meal results collected across Australia from more than 12,000 blood meals from 22 species. To assess mosquito-vertebrate associations and identify mosquitoes on a spectrum of generalist or specialist feeders, we analysed blood meal data in two ways; first using a novel odds ratio analysis, and secondly by calculating Shannon diversity scores. We find that each mosquito species had a unique feeding association with different vertebrates, suggesting species-specific feeding patterns. Broadly, mosquito species could be grouped broadly into those that were primarily ornithophilic and those that fed more often on livestock. Aggregated feeding patterns observed across Australia were not explained by intrinsic variables such as mosquito genetics or larval habitats. We discuss the implications for disease transmission by vector mosquito species classified as generalist-feeders (such as *Aedes vigilax* and *Culex annulirostris*), or specialists (such as *Aedes aegypti*) in light of potential influences on mosquito host choice. Overall, we find that whilst existing blood meal studies in Australia are useful for investigating mosquito feeding patterns, standardisation of blood meal study methodologies and analyses, including the incorporation of vertebrate surveys, would improve predictions of the impact of vector-host interactions on disease ecology. Our analysis can also be used as a framework to explore mosquito-vertebrate associations, in which host availability data is unavailable, in other global systems.

## Introduction

Mosquitoes are the most important disease vector globally, responsible for infecting millions of people and animals annually with pathogens that influence human health, livestock and economic trade and wildlife biodiversity [1]. Mosquitoes comprise a broad taxonomic group with more than 3000 species recognised across 40 genera [2], but not all species are involved in pathogen transmission. Pathogen transmission requires a mosquito to take a bloodmeal from a source host and then to subsequently feed on a recipient host. Understanding the feeding patterns of mosquitoes can inform disease management strategies (such as targeted vector control to reduce vector-host contact), and can contribute to models forecasting future disease risk in human and animal populations [3].

Mosquito host choice is complex; both intrinsic and extrinsic factors can influence feeding preference [3, 4]. Intrinsic variables can include genetics, whereby individuals are more likely to feed on the same host as previous generations [5, 6], and the nutritional state of the mosquito, with nutrition-poor individuals being more likely to feed on non-preferred hosts [4]. Extrinsic host-seeking behaviour is predominantly guided by detection of heat and carbon dioxide (CO2), and is also affected by host abundance, biomass, various odorants and chemicals that are released by hosts, and host defensive behaviour [7-13]. Other extrinsic factors may include climatic variables such as relative humidity, along with habitat characteristics that determine availability and diversity of hosts [12, 14]. In addition to these broad intrinsic and extrinsic variables, evidence suggests that mosquitoes may adjust their host-seeking behaviours based on positive and negative experiences, in essence, adapting feeding choices according to their individual circumstances [4].

Mosquito-host relationships in Australia are largely understudied. The island biogeography of Australia and its varied climatic zones and bioregions promote a unique endemic biodiversity for mosquito and vertebrate host species. Along with a high diversity of native marsupials, placental mammals and birds in Australia, there are more than 300 species of mosquitoes described, many of which are unique to the continent [15]. The interactions between these populations of mosquitoes and vertebrate hosts across different climatic zones provide opportunities for maintenance and emergence of mosquito-borne pathogens. The transmission of numerous medically-important arboviruses has been documented in Australia to date, including dengue (DENV), Ross River (RRV), Murray valley encephalitis (MVEV), Barmah Forest (BFV) and Kunjin [16] viruses.

Taking into account the complexity of contributing factors, critical analysis of the feeding patterns of mosquitoes may represent an important approach to explore disease risks for both human and animal populations. This study aims to synthesise existing literature describing blood meal studies in Australia, specifically assessing the most likely mosquito-host associations, and the diversity of feeding patterns for common mosquito species. In light of these feeding patterns, we discuss broad implications for disease ecology.

## Methods

### Data collection

Original research articles were systematically searched by using the following search terms in different combination across five search engines (Web of Science, ProQuest, Science Direct, PubMed and Google Scholar): ‘bloodmeal*’, ‘blood meal’, blood-meal’, ‘feeding’, ‘habit’, ‘pattern*’, ‘preference*’, ‘interaction*’, ‘mosquito*’, ‘vector*’, ‘vector-host’, ‘host*’, ‘vertebrate*’, ‘animal*’ and ‘Australia’. The asterisk (*) operator was used as a wildcard to search for all possible variations of keywords. We then manually searched the reference lists of papers to identify additional relevant articles. Papers were included in this review if they were original peer-reviewed research articles, undertaken on mainland Australia (e.g. [17] was undertaken in Badu Island in the Torres Strait and thus excluded), analysed field collected mosquitoes that had fed under natural conditions on free-living vertebrates (e.g. [18] used tethered animal baits and was excluded), and assessed at least 3 potential vertebrate species (e.g. [19] only tested for a single flying fox species, and was excluded).

The following information was extracted from identified articles: the geographic area in which the study took place (including site location, bioregion of each site (as defined by Thackway and Cresswell [20]); the mosquito collection method used (including the year, month and collection method/trap type); and the methods used to determine the vertebrate origin of blood meals (including the vertebrate species investigated, the source of vertebrate reference samples, and laboratory technique). Additional notes were made on stated limitations (if any) of each paper and whether data on vertebrate abundance and diversity was included. A database of blood meal results was populated and is reported in Supplementary Table 2.

### Data analysis

#### Mosquito-vertebrate associations

Odds ratios were used to calculate the direction (positive or negative) and strength of associations in the database between each mosquito species and vertebrate taxon. For this analysis, the blood meal origin data compiled from the literature were aggregated such that each vertebrate species was grouped into the broader taxonomic groups of Humans, Carnivores (cats, dogs and foxes), Aves (all birds), Diprodontia (all possum and kangaroo species), Artiodactyla (cows, sheep, pigs and goats) and Perisodactyl (horses). Flying fox [21], rodent [22, 23] and rabbit [22, 24] species were excluded from this analysis, as the sample size of blood meals from these species was too small (either one or two studies, with these species comprising less than 9% of blood meal origins within each). To be included in the analysis, mosquito species needed to meet all minimum data criteria of i) their blood meal origins being reported more than twice in the literature, and ii) having an arbitrary minimum of 35 blood meals identified.

Log odds ratios were calculated between each mosquito and vertebrate taxon using two by two feeding frequency tables, derived from the raw data (Supplementary Table 1). Positive log odds ratios indicate a positive feeding association between a given mosquito species and vertebrate host, whereby there is a higher likelihood than random chance that a blood meal of that mosquito species would originate from the given vertebrate taxon. The greater the log odds ratio, the stronger the feeding association. Conversely, a negative log odds ratio suggested a negative feeding association, whereby there is a lower likelihood that a blood meal from the mosquito species would originate from the given vertebrate taxon. Log odds ratios close to 0 indicate no association between the mosquito species and vertebrate taxon.

The log odds ratios were plotted in a heatmap chart and sorted using hierarchical clustering. The clustering grouped mosquitoes with similar feeding patterns together by similarity in log odds ratio across all vertebrate taxa. All calculations and graphs were generated using R software, with packages *gplots* and *RColorBrewer* [25] with modified script from Raschka [26].

#### Mosquito feeding diversity

We used the Shannon diversity index to place mosquito species on a spectrum between generalist or specialist feeders. The inclusion criteria for this analysis was that each mosquito species needed to have fed on greater than three vertebrate species and had to have a minimum number of 10 blood meals analysed. Vertebrates were not aggregated by taxonomic group in this analysis, but remained at the level reported in the literature (mostly as species but, for the case of birds, several studies reported as class). A total of 15 vertebrate species were included in this analysis as blood meal origins, and 13,934 blood meals from 21/41 mosquito species met the criteria (Supplementary Table 2).

Shannon diversity indices were calculated for each mosquito species [27] and expressed as an h-index. A higher h-index is associated with a greater feeding diversity, as it suggests a mosquito species has fed on a greater number of vertebrate species and/or feeds evenly across vertebrates. Conversely a lower h-index suggests mosquito species have a low feeding diversity, and are associated with feeding on fewer vertebrate species and/or a greater number of feeds on a small number of vertebrates. Within this dataset, we categorised an h-index in the top quartile as ‘high feeding diversity’, whilst an h-index in the lowest quartile was considered a ‘low feeding diversity’. Shannon diversity indices were calculated in Excel.

## Results

### Characteristics of the selected studies

We identified ten papers that met the search criteria, comprising 14,044 mosquito blood meals across 48 mosquito species. Study characteristics and methodologies are summarised in Table 1. These studies took place at 32 sites across 14 bioregions, in all mainland states and territories in Australia (Figure 1). The selected studies were undertaken over a 62 year-period, from 1954 to 2016.

**Table 1:** Location and methods for mosquito collection and blood meal analysis for the studies included in this review

| Study (Year) [Reference] | Site name (state) | Site habitat type | Collection months (years) | Mosquito collection method | Number of blood meals (& species) analysed | Blood meal analysis method | Vertebrate species tested |
| --- | --- | --- | --- | --- | --- | --- | --- |
| Flies *et al.* (2016) [23] | Adelaide Hills, Adelaide City, Murray River valley (South Australia) | Urban and rural | August – May (2006-2015) | EVS + CO<sub>2</sub> | 199 (8) | Cytochrome b PCR using vertebrate, avian and mammalian primers, gene sequencing | Dog, cat, cow, sheep, pig, chicken, brushtail and ringtail possum, red kangaroo, koala, rainbow lorikeet, galah, Australian magpie |
| Hall-Mendelin *et al.* (2012) [34] | (Queensland), (Northern Territory) | Rural | February, March, October (2002-2006) | CDC + CO<sub>2</sub> +/- octenol | 1,128 (1) | Double-antibody ELISA modified from | Horse, rat, human, dog, cat, bird, kangaroo, cow, pig |
| Johansen *et al.* (2009) [24] | 32 sites (Western Australia) | Urban and rural | All year round (1993-2004) | EVS + CO<sub>2</sub> | 2,606 (29) | Double-antibody ELISA modified from | Cow, sheep, goat, pig, rabbit, horse, donkey, human, cat, fox, dog, brushtail possum, quokka, Western grey kangaroo, mouse, chicken, duck |
| Jansen *et al.* (2009) [22] | Cairns (Queensland), Brisbane (Queensland), Newcastle (New South Wales), Sydney (New South Wales) | Urban | All year round (2005-2008) | CDC, EVS + CO<sub>2</sub>; Unbaited BG sentinel traps | 1,180 (15) | Indirect ELISA; PCR using avian- and mammalian-specific primers, gene sequencing | Horse, rabbit, rat, human, dog, chicken, cat, bird, kangaroo, cow, pig, 50 wild bird species |

| Study<br>(Year)<br>[Reference] | Site name (state) | Site habitat type | Collection<br>months<br>(years) | Mosquito<br>collection<br>method | Number of<br>blood<br>meals (&<br>species)<br>analysed | Blood meal<br>analysis method | Vertebrate species tested |
| --- | --- | --- | --- | --- | --- | --- | --- |
| Kay *et al.*<br>(2007) [21] | Brisbane<br>(Queensland) | Urban | September-April (2000-2001) | CDC + CO<sub>2</sub> +/-<br>OCT | 865 (10) | Gel diffusion<br>immunoassay | Bird, kangaroo, cat, dog,<br>horse, human, brushtail<br>possum, flying fox species |
| Frances *et al.*<br>(2004) [32] | Shoalwater Bay<br>(Queensland) | Rural | All year round<br>(1998-2000) | EVS + CO<sub>2</sub> | 763 (15) | Gel diffusion<br>immunoassay | Human, kangaroo, bird, dog,<br>horse, cow |
| van den<br>Hurk *et al.*<br>(2003) [33] | Cape York, Gulf<br>Plains<br>(Queensland) | Rural | January-May<br>(1995-2001) | CDC + CO<sub>2</sub> +/-<br>OCT | 2,582 (15) | Gel diffusion<br>immunoassay | Horse, rabbit, rat, human,<br>dog, chicken, cat, bird,<br>kangaroo, cow, pig |
| Muller *et al.*<br>(1981) [29] | Beatrice Hill<br>(Northern<br>Territory) | Rural | All year round<br>(1974-1976) | Light trap (not<br>specified) and<br>vehicle<br>mounted trap | 1,628 (18) | Preciptin test | Cow, horse, dog, human,<br>marsupial, chicken |
| Kay *et al.*<br>(1979) [30] | Charleville<br>(Queensland) | Rural | February<br>(1976) | Aspiration of<br>resting sites | 5,431 (16) | Preciptin test | Reptile, amphibian, bird,<br>pig, dog, cat, man, cow,<br>horse, rodent, bat, marsupial,<br>carnivore |
| Lee *et al.*<br>(1954) [31] | Moree, Hornsby<br>(New South<br>Wales), Texas<br>(Queensland),<br>Golburn Valley<br>(Victoria),<br>Canberra<br>(Australian<br>Capital Territory) | Mostly rural | December-January<br>(1951-1952) | Aspiration of<br>resting sites | 1,231 (15) | Preciptin test | Human, chicken, cow, dog,<br>rabbit, horse, marsupial<br>(unspecified) |
| **Total** |  |  |  |  | **14,044 (48)** |  |  |

Abbreviations: Encephalitis Vector Survey (EVS); Carbon Dioxide (CO<sub>2</sub>); Centers for Disease Control (CDC); BioGent (BG); Polymerase chain reaction (PCR); Enzyme-linked immunosorbent assay (ELISA); Octane (OCT)

**Figure 1:**
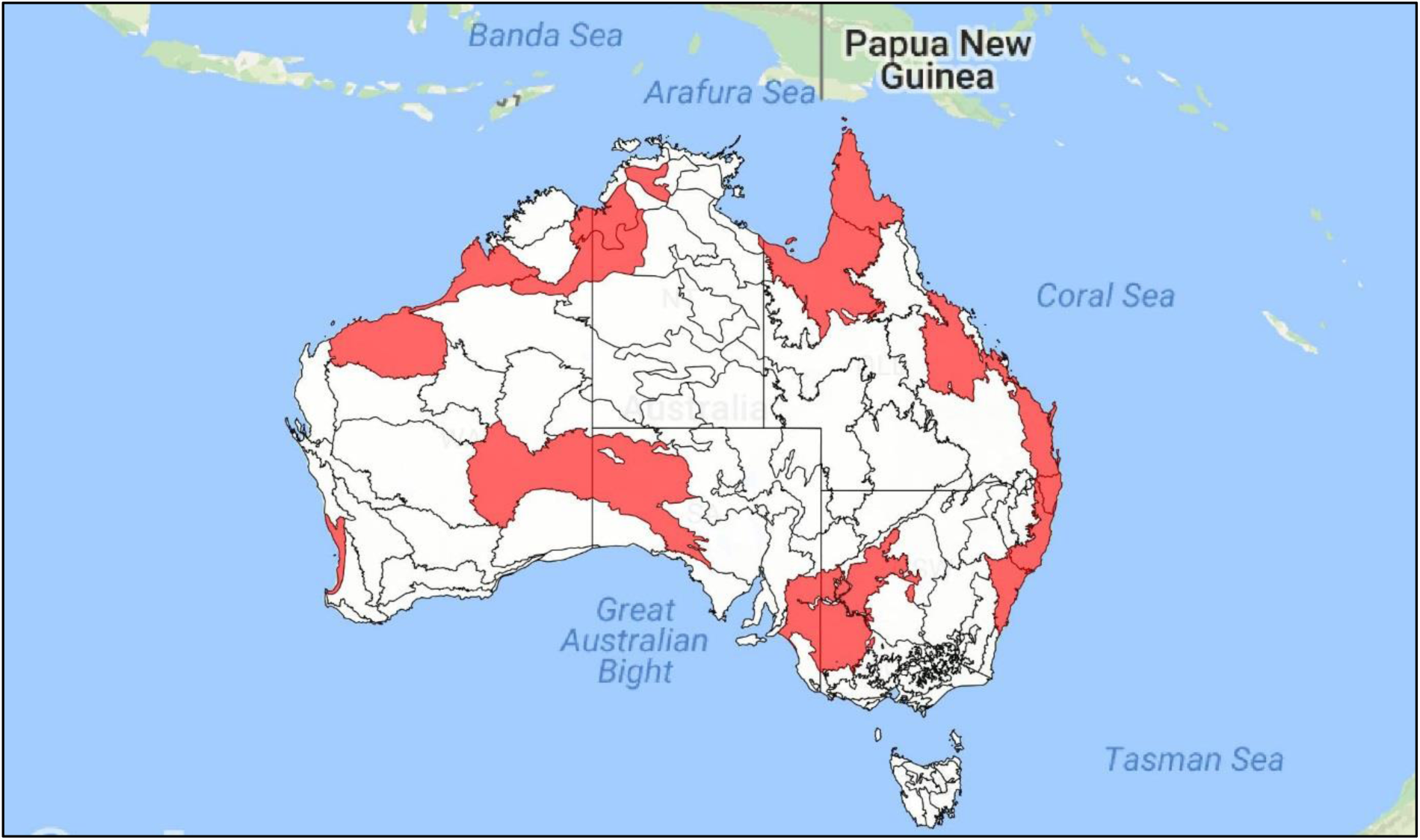
Bioregions in which blood meal studies took place (indicated in red) across Australia

To collect blood-fed mosquitoes, most studies (n = 8) used Centers for Disease Control (CDC) CO2-baited miniature light traps [28], supplemented with 1-octen-3-ol in some cases (Table 1). Other methods included unbaited BioGent® (BG) sentinel traps [22] and aspiration of resting sites [21, 29, 30]. One study [29] also used vehicle-mounted traps. To analyse blood meals, early studies employed preciptin tests [29-31] and serological gel diffusion techniques [32, 33] (Table 1). More recent studies adopted enzyme-linked immunosorbent assay (ELISA) and various molecular techniques including polymerase chain reaction (PCR) and gene sequencing [22-24, 34]. Vertebrate reference sources most commonly employed in immunoassays and gel diffusion techniques were commercially-available anti-sera and included horse, rabbit, rat, dog, chicken, cat, bird, kangaroo, cow and pig. Molecular studies which included wildlife used vertebrate references provided through wildlife hospitals, zoos and roadkill. These studies also included DNA sequence data available on GenBank.

### Mosquito-vertebrate associations

Of the 10 studies of blood meal origins, data on 41 mosquito species were reported, and data on 12 of these met the criteria to be included in analysis; *Aedes normanensis, Ae. notoscriptus, Ae. procax, Ae. vigilax, Anopheles annulipies, An. bancroftii, Coquillettidia linealis, Cq. xanthogaster, Culex annulirostris, Cx. sitiens, Cx. quinquefasciatus* and *Mansonia uniformis* (Figure 2). All species, except *Ae. procax*, showed significant positive associations with at least one vertebrate host. The strongest positive log odds ratio was between *Cx. annulirostris* and the Diprodontia taxa (possums and kangaroos; log odds ratio (LOR) = 2.77), followed by *Ma. uniformis* and humans (LOR = 2.2). All mosquito species except *Ae. vigilax* and *Cq. xanthogaster* had strong negative association with at least one vertebrate taxon. The strongest negative log odds ratio was between *Cx. quinquefasciatus* and Diprodontia (LOR = -3.8), followed by *Ae. notoscriptus* and Artiodactyla (cows, sheep, pigs and goats; LOR = -2.8).

**Figure 2.**
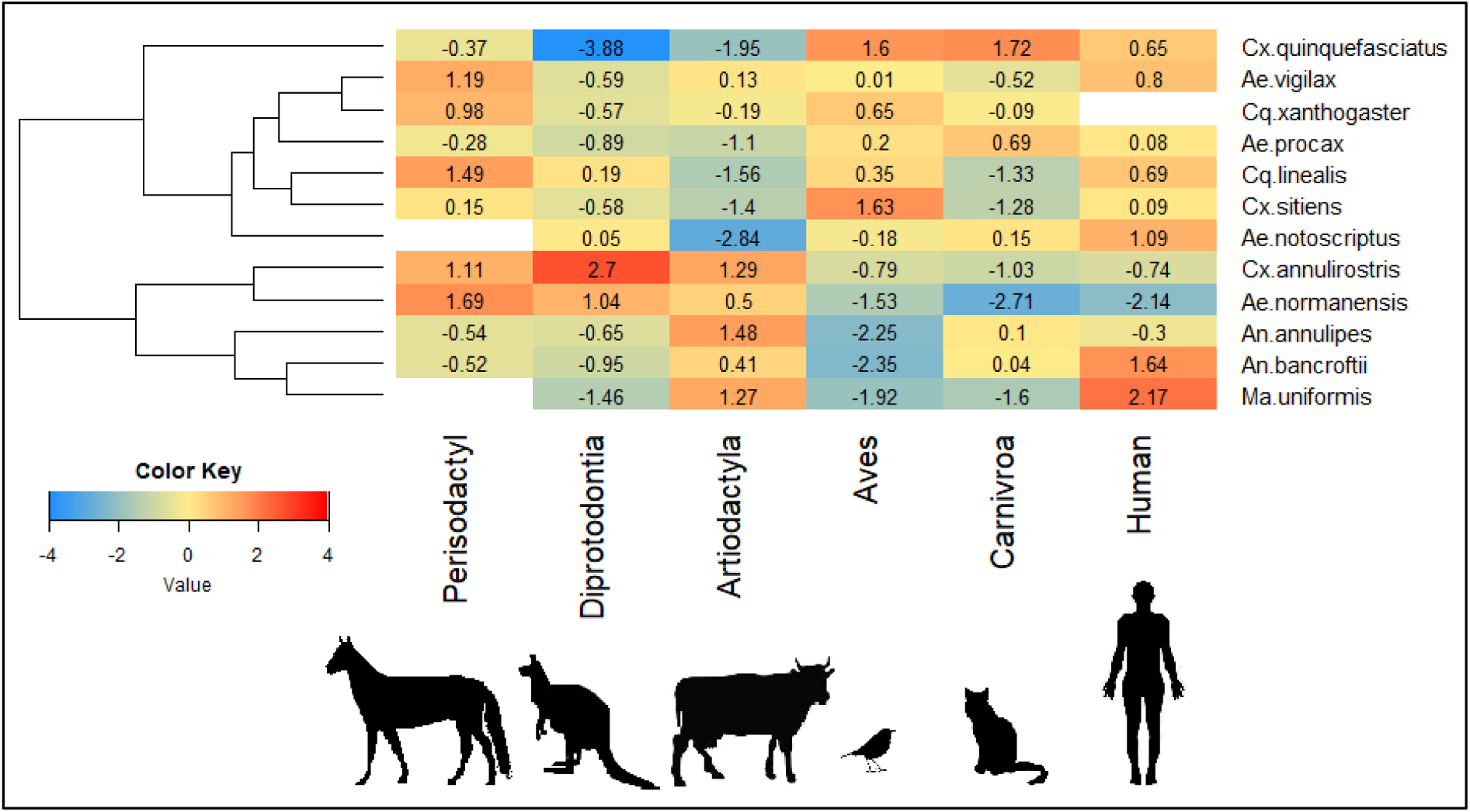
Feeding associations between Australian mosquito species and vertebrate taxa. Log odds ratio for mosquito species (right hand side) indicate feeding likelihood on vertebrate taxa (bottom). Each are sorted by hierarchal cluster (left) according to similarities in feeding ratios between mosquito species.

The mosquito species clustered together in two broad groups. The first cluster group consisted of seven mosquito species (*Cx. quinquefasciatus, Ae. vigilax, Cq. xanthogaster, Ae. procax, Cq. linealis, Cx. Sitiens* and *Ae. notoscriptus*), of which most shared a negative association with the Artiodactyla (6 of the 7 species) and Diprodontia vertebrates (5/7), and a positive association with Humans (7/7) and Aves (6/7). Within this cluster, *Ae. vigilax* and *Cq. xanthogaster* were in the same clade and shared a strong positive association with Perisodactyla. *Coquillettidia linealis* and *Cx. sitiens* also shared a clade and strong negative association with both Aritodactyl and Carnivora.

The second major cluster group consisted of five mosquito species (*Cx. annulirostris, Ae. normanesis, An. annulipies, An. bancroftii* and *Ma. uniformis*). These species all shared a positive association with Aritodactyla and a negative association with Aves. *Culex annulirostris* and *Ae. normanesis* were on the same clade and shared a strong positive association with Perisodactyl and Diprodontia. Although they both also had a negative association with Aves, Carnivora and Humans, this was strongest only for *Ae. normanensis*. *Anopheles annulipies* and *Ma. uniformis* were on a single clade and were both had high associations with humans.

### Mosquito feeding diversity

Twenty-two mosquito species met the criteria for inclusion in the Shannon diversity analysis analysis (Figure 3), comprising 12,424 individual blood meals in total. The median h-index reported across all species was 1.40, and the mean was 1.34. Low feeding diversity (h-index = <0.99) was observed in five mosquito species; of which *Ae. aegypti* had the lowest diveristy (h-index = 0.72). High feeding diversity (h-index = >1.64) was reported in five mosquito species; of which *Ae. vigilax* had the highest diversity (h-index = 2.17).

**Figure 3:**
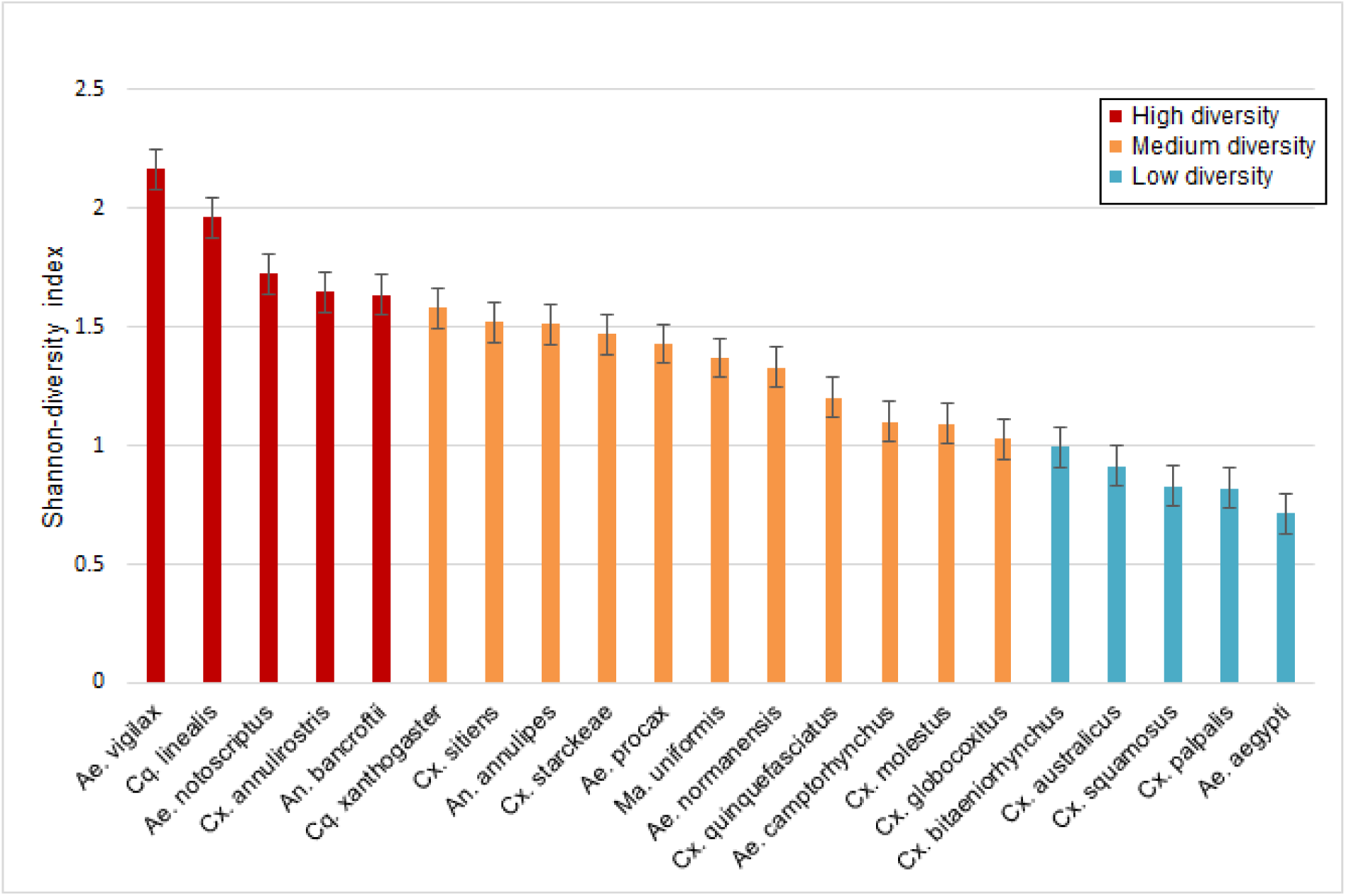
Shannon diversity (h-index) of blood meal origins for Australian mosquitoes, error bars represent the standard error for all measures

## Discussion

Our analysis found that each mosquito species had a unique feeding association with different vertebrates, suggesting species-specific feeding patterns. The hierarchical clustering from the odds ratio analysis sorted mosquitoes in two broad groups; mosquitoes that either had a positive association with birds (Aves) and negative association with livestock (Artiodactlya), or vice-versa. Interpreting the feeding patterns of these particular mosquito species is important, given that at least half of these mosquitoes have been found to be competent vectors for notifiable arboviruses in Australia [35-40], whilst the other half have been demonstrated to carry some viruses, though their ability to transmit them hasn’t been fully investigated [41-43].

Intrinsic drivers of mosquito host choices (such as genetics, larval ecology and dispersal) did not explain feeding patterns in this analysis. Specifically mosquito species did not group together by taxonomic relatedness (e.g. genus). Studies examining the effect of genetics on mosquito host choices have found that offspring are more likely to feed on the same host as previous generations [5, 6]. However, this has only been demonstrated within-species and is unlikely to be important between species belonging to same genus, particularly since potential for rapid evolution (due to short generation span) likely reduces the influence that taxonomic relatedness may have on mosquito feeding host behaviour. Another intrinsic factor, mosquito larval ecology, may partially explain some clustering. For example, *Ae. normanensis* and *Cx. annulirostris* grouped together and larvae of both typically inhabit inland freshwater; similarly, the larval habitat of both *An. bancroftii* and *Ma. uniformis* is freshwater swamps. Although this pattern did not explain all clusters, it implies that local environmental influences, at least partially, drive mosquito host choice. This is perhaps not surprising when considering potential limitations on dispersal from larval habitats for various mosquito species. For example, *Ae. vigilax* is recognised as having large dispersal capability, being found more than 50 km from potential saltwater larval habitats, albeit likely wind-assisted in some cases [44]. This high dispersal potential suggests that *Ae. vigilax* can move readily between locations, allowing feeding on a diversity of vertebrate taxa, as reflected in our feeding diversity analysis. As such, whilst genetics and larval habitats may be important within species on a local scale, they do not explain the aggregated feeding patterns observed across Australia.

Extrinsic variables, such as species abundance and diversity, explain in part some of the feeding associations in this analysis, but not all. Mosquitoes have complex interactions with their environment. Thus, factors broader than vertebrate abundance alone are important to consider for mosquito feeding patterns. For example, mosquito flying/resting height has been linked to host feeding patterns [45-48]. In two Australian studies, more *Cx. sitiens* and *Cx. quiquefasciatus* were caught in traps set at least 8m off the ground, whilst a higher abundance of *Ae. vigilax* were found in traps 1.5m off the ground, for the same locations [47, 48]. In our meta-analysis, *Cx. sitiens* and *Cx. quinquefasciatus* had strong positive associations with blood meals originating from tree dwelling bird species (i.e. Australasian figbirds *Sphecotheres vieilloti*, Common myna *Sturnus tristis* and Helmeted friarbirds *Philemon buceroides* [22]), whilst *Ae. vigilax* had the strongest positive associations with ground dwelling species (horses and humans). This could suggest that whilst overall vertebrate abundance within a given environment can influence the availability of a particular host, mosquitoes are highly mobile and may seek a blood meals across ecological niches a given habitat. As such different mosquito species can exhibit different feeding patterns despite being exposed to the same vertebrate abundance and climate conditions in the same habitat.

In the odds ratio analysis, *Cx. annulirostris* exhibited an unexpected feeding pattern. This species is considered an important vector for medically-important arboviruses [36, 49, 50], however the meta-analyses which included more than 5700 blood meals, found that *Cx. annulirostris* had only a weak feeding association with humans (LOR = -0.74). This is consistent with an early field study assessing mosquito feeding preferences using live baits, in which *Cx. annulirostris* preferred cows, pigs and dogs more than humans [30]. The Shannon diversity analysis, along with other studies, have identified *Cx. annulirostris* as a generalist feeder with plastic feeding patterns that may shift temporally or spatially [51, 52]. This knowledge, in combination with the widespread distribution of *Cx. annulirostris* across Australia, suggests that localised studies of *Cx annulirostris* feeding are required to assess the role the species plays in disease transmission for which it is theoretically an important vector.

In addition to *Cx. annulirostris, Aedes vigilax* and *Ae. notoscriptus* were identified as generalists due to their high diversity scores in the Shannon diversity analysis. International studies [53-55] suggest that generalist feeders are capable of playing a role as bridge vectors due to their ability to acquire pathogens from animal hosts, and subsequently transmitting the pathogen to humans. Bridge vectors are particularly important for enzootic amplification of arboviruses and are often associated with outbreaks [53]. For the species identified in this analysis as generalists, they have been demonstrated to be competent vectors of zoonotic arboviruses in Australia [35, 56-58], and as such should be closely monitored to reduce transmission between vectors and humans.

The disease ecology associated with specialist feeders is also important to consider. Here we identified *Ae. aegypti* as having the lowest feeding diversity, indicating the species as a specialist feeder. Indeed, more than 70% of the blood meals originated from humans. The anthropophilic feeding observed in *Ae. aegypti* is similar to that reported in international studies, where 80 to 99% of all blood meals are human in origin [59, 60]. This feeding pattern for *Ae. aegypti* is consistent with its role as an important vector of several arboviruses which are transmitted between humans without an animal reservoir, including dengue, Zika and chikungunya viruses. Interestingly, although the importance of *Ae. aegypti* is recognised, *Ae. aegypti* is sometimes under laboratory conditions the species has been observed to demonstrate relatively poor transmission rates for DENV, when compared to other mosquito species [61, 62]. In this case, being a specialist feeder, preferring mainly humans, is what determines the status of *Ae. aegypti* as an important disease vector, rather than its competence [63].

### Future directions

An absence of data on host availability in the regions where mosquitoes were collected limits inferences on host preference specifically. Of the blood meal studies reviewed here, only one considered host abundance [21]. That study assessed abundance through a local resident survey on the number of pets, people and estimated number of possums in the vicinity, adding confidence to the interpretation of vector-feeding patterns. Such collection of host ecology data in conjunction with blood fed mosquitoes can be considerably labour intensive; however, it provides a more thorough assessment of how host abundance and biomass may influence observed mosquito feeding patterns and informs the selection of appropriate reference samples against which to compare blood meals in the laboratory. Where data cannot be collected in conjunction with blood fed mosquitoes, alternative sources such as the Atlas of Living Australia and the Global Biodiversity Information Facility (GBIF) may offer a suitable proxy. This methodology has been adopted successfully in international blood meal studies [64, 65] and could be beneficial for future investigations.

Although a range of reference vertebrates were often included in Australian blood meal studies, they were rarely a true representation of the vertebrates available to mosquitoes for feeding. At present there are large gaps in understanding the role of cryptic, migratory or smaller mammalian species in mosquito feeding patterns. For example, only two studies included rabbits [22, 24] and rodents [22, 23] in their analysis. Despite their small size, rabbits and rodents were identified to be the origin of blood meals for *Cx. sitiens* and *Cq. linealis* [22]. Mosquito-rodent associations have also been identified in the literature, where by at least 27% of mice were seropositive to RRV [58, 66]. It is therefore important that, despite small body size, rats and rodents are included in future investigations of mosquito blood meals.

### Conclusion

Improved understanding of mosquito feeding patterns can lead to better management and risk predictions for medically important arboviruses. Here we find that of the Australian mosquito species tested, each had a unique feeding pattern; however, the particular specialist or generalist feeding patterns of mosquito species could be a key determinant of the risk they pose for human disease. These patterns, and the resulting human disease risk, are likely influenced by a suite of intrinsic and extrinsic variables. Broader ecological considerations alongside these feeding patterns could be useful for the interpretation of these complex biological systems, but at present data available to do this is limited. Future studies should utilise multidisciplinary approaches to collect data on vertebrate communities in parallel with mosquito communities. More data from both top-down (broad assessments of blood meals) and bottom-up approaches (specialised host choice experiments) are needed in conjunction with modelling techniques to bring these data together for meaningful interpretation of arbovirus transmission risk in Australia.

---

**Supplementary Table 1:** Derivation of 2×2 contingency tables for mosquito feeding preferences

|  | Focal host taxon | All other hosts | Totals |
| --- | --- | --- | --- |
| Focal mosquito species | $a$ | $A-a$ | $A$ |
| All other mosquito species | $B-a$ | $N-A-B+a$ | $N-A$ |
| Totals | $B$ | $N-B$ | $N$ |

$a$ Number of records of the focal mosquito species feeding on the focal host taxon $A$ Total number of records for the focal mosquito species $B$ Total number of records for the focal host taxon $N$ Overall number of records Values in grey shaded cells are obtained by subtraction

**Supplementary Table 2:**
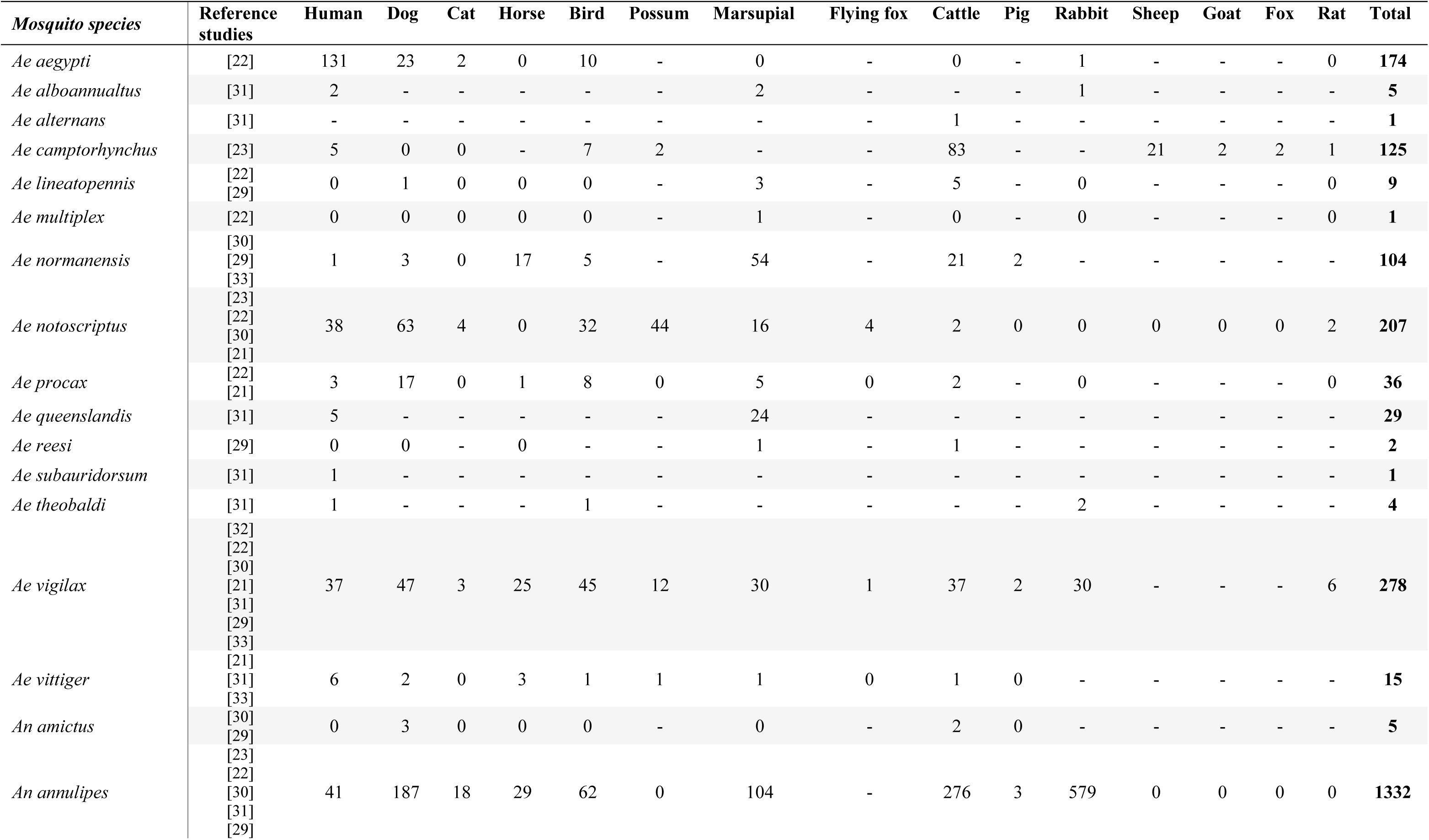
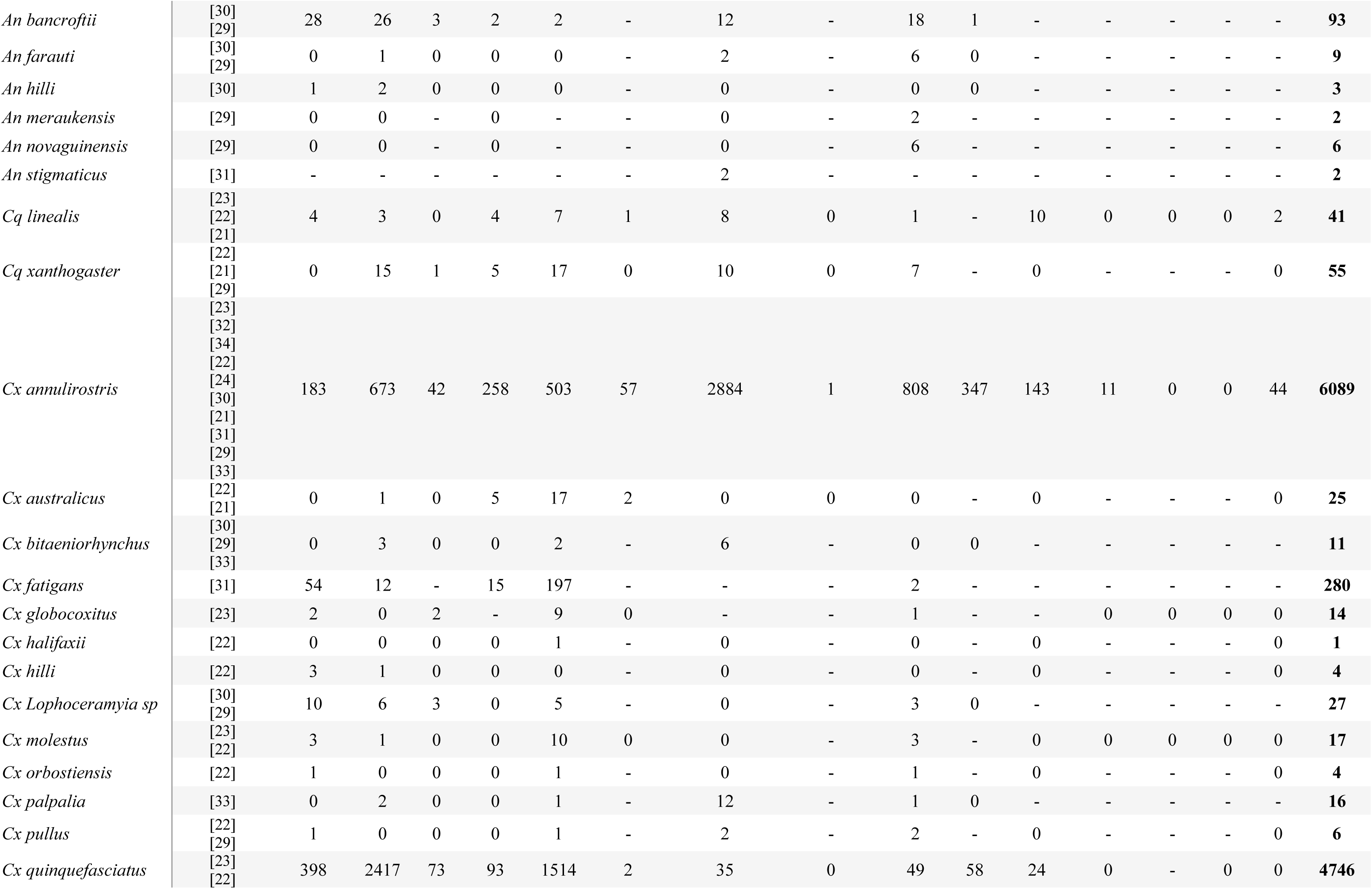
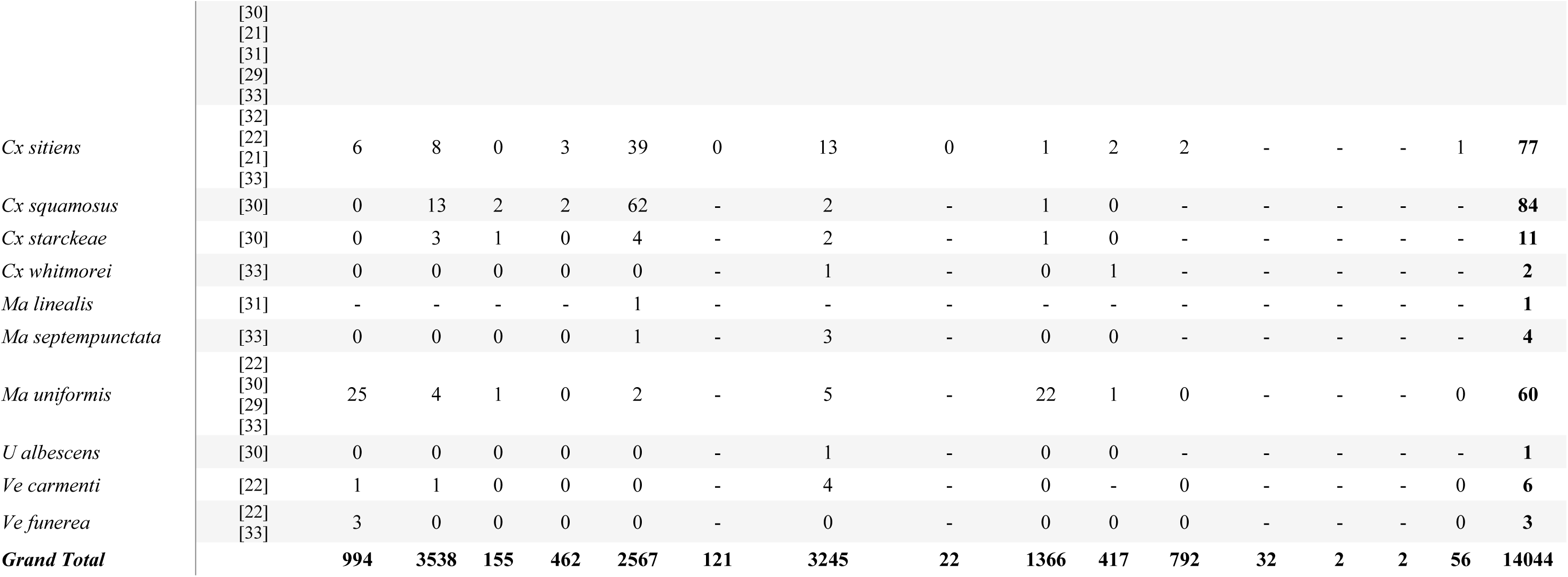
Reported blood meal results for Australian mosquito species. Numbers in the columns representing number of blood meals for each vertebrate.

